## Supplemental Figures for "MicroRNA-26b protects against MASH development in mice and can be efficiently targeted with lipid nanoparticles"

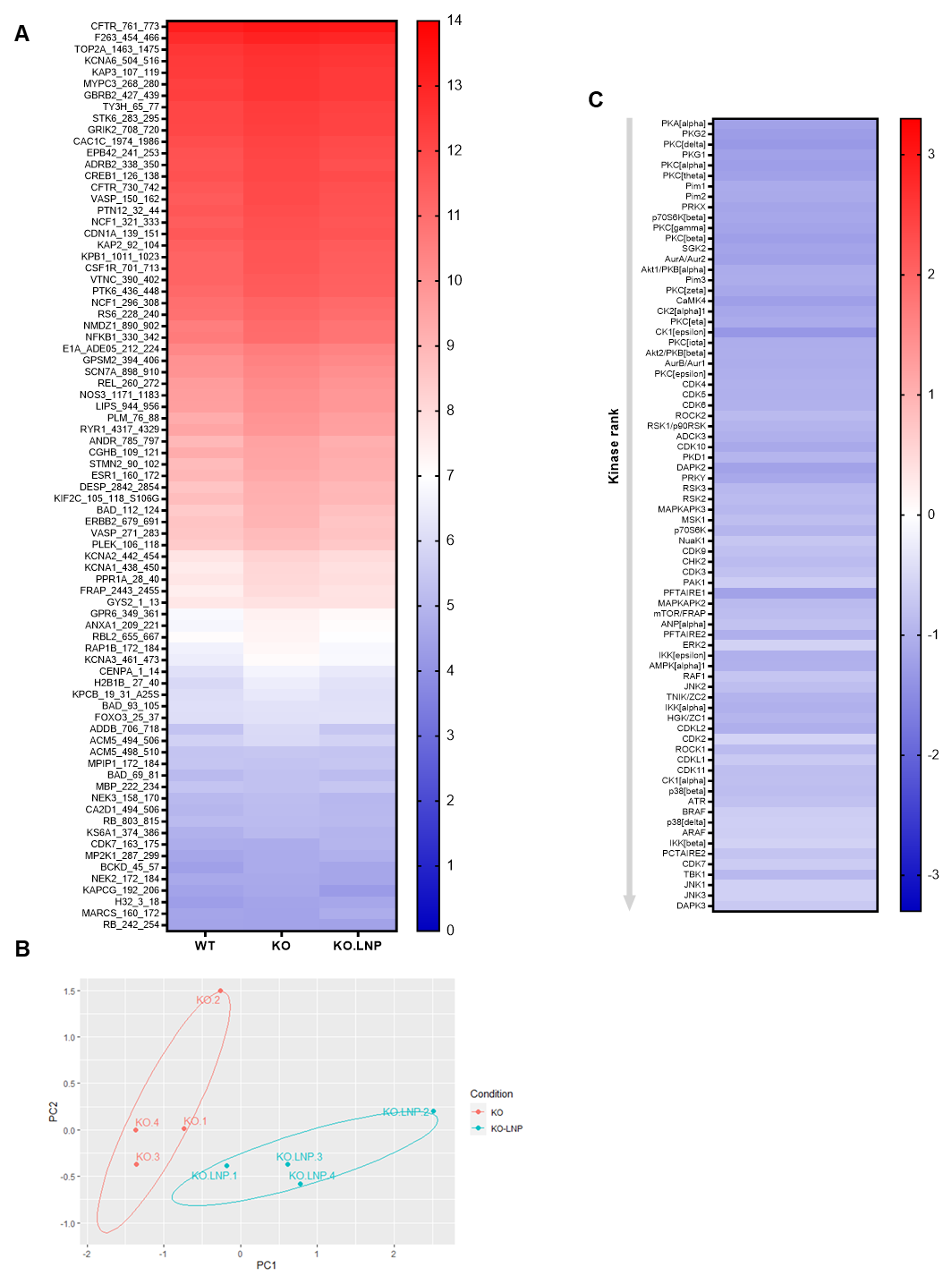

**Supplemental Figure 1. mLNP treatment rescues the inflammatory kinase activity effect of *miR-26b* knockout.**

(**A**) Heatmap demonstrating the level of peptide phosphorylation (numbers behind peptides indicate exact amino-acids that are spotted on the STK array). Red color reflects a high degree of phosphorylation, while blue color represents a low degree of phosphorylation (average of n=4 is shown). (**B**) Principal component analysis (PCA) of phosphorylated peptides from STK array (n=4) of liver lysates from mLNP-treated *Apoe^-/-^Mir26b^-/-^* mice (KO.LNP) or *Apoe^-/-^Mir26b^-/-^* mice (KO) mice**.** (**C**) The heatmap of significantly changed kinases is ranked based on Median Final Score (cut-off value of 1.2), STK array performed on liver lysates from mLNP treated *Apoe^-/-^Mir26b^-/-^* mice (KO.LNP) compared to *Apoe^-/-^Mir26b^-/-^* mice (KO) mice**.** Color corresponds to the Median Kinase Statistic, which represents effect size and directionality (blue = decreased activity in KO.LNP vs. KO mice).

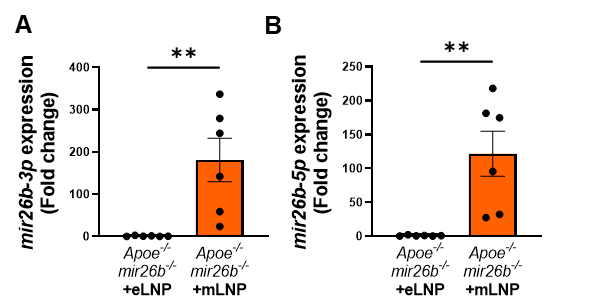

**Supplemental Figure 2. mLNP treatment overexpresses miR26b-3p and -5p in murine livers.**

(**A-B**) Gene expression analysis of (**A**) *mir26b-3p* and (**B**) *mir26b-3p* in livers from mice after 4-week WTD with simultaneous injection with either empty LNPs as vehicle control (eLNP) or LNPs containing miR-26b mimics (mLNP) every 3 days. ***p*<0.01. *n*=6 animals per group.
